# Micapipe: A Pipeline for Multimodal Neuroimaging and Connectome Analysis

**DOI:** 10.1101/2022.01.31.478189

**Authors:** Raúl R. Cruces, Jessica Royer, Peer Herholz, Sara Larivière, Reinder Vos de Wael, Casey Paquola, Oualid Benkarim, Bo-yong Park, Janie Degré-Pelletier, Mark Nelson, Jordan DeKraker, Christine Tardif, Jean-Baptiste Poline, Luis Concha, Boris C. Bernhardt

## Abstract

Multimodal magnetic resonance imaging (MRI) has accelerated human neuroscience by fostering the analysis of brain structure, function, and connectivity across multiple scales and in living brains. The richness and complexity of multimodal neuroimaging, however, demands processing methods to integrate information across modalities and different spatial scales. Here, we present *micapipe*, an open processing pipeline for BIDS-conform multimodal MRI datasets. *micapipe* can generate i) structural connectomes derived from diffusion tractography, ii) functional connectomes derived from resting-state signal correlations, iii) geodesic distance matrices that quantify cortico-cortical proximity, and iv) microstructural profile covariance matrices that assess inter-regional similarity in cortical myelin proxies. These matrices are routinely generated across established 18 cortical parcellations (100-1000 parcels), in addition to subcortical and cerebellar parcellations. Results are represented on three different surface spaces (native, conte69, fsaverage5), and outputs are BIDS-conform. Processed outputs can be quality controlled at the individual and group level. *micapipe* was tested on several datasets and is available at https://github.com/MICA-MNI/micapipe, documented at https://micapipe.readthedocs.io/, and containerized as a BIDS App http://bids-apps.neuroimaging.io/apps/. We hope that micapipe will foster robust and integrative studies of human brain microstructure, morphology, and connectivity.

## 1. Introduction

The human brain is a highly complex network organized across multiple spatial and temporal scales (Betzel and Bassett 2017). Neuroimaging, and in particular magnetic resonance imaging (MRI), provides versatile contrasts sensitive to the brain’s microstructure, connectivity, and function, offering a window into its organization in living humans (Turner, 2019; Larivière et al., 2019; van den Heuvel et al., 2019; Van Essen et al., 2013).

Recent years have witnessed multiple neuroimaging data acquisition efforts (Gordon et al., 2017; Royer et al., 2021; Van Essen et al., 2012) as well as initiatives for open data sharing to promote transparency and reproducibility (Milham et al., 2018). These initiatives offer researchers the ability to interrogate brain structure and function in thousands of individuals across multiple sites from around the world. In addition, a variety of processing pipelines has previously been developed. These include tools for the automated analysis of cortical/subcortical morphology based on T1-weighted MRI (Fischl, 2012; Kim et al., 2005; Patenaude et al., 2011), approaches for the analysis of myelin-sensitive MRI contrasts to assess brain microstructure (Paquola et al., 2019b; Glasser and Van Essen, 2011; Waehnert et al., 2016), the study of intrinsic brain function and functional connectivity via resting-state functional MRI, rs-fMRI (Biswal et al., 2010; Craddock et al., 2013; Esteban et al., 2019), and analysis of structural connectivity inferred via diffusion MRI tractography (Cieslak et al., 2021; Daducci et al., 2012; Tournier et al., 2019). Individually, ongoing advances in MRI modelling approaches result in increasing biological validity (Craddock et al., 2015; Jbabdi et al., 2007; Mars et al., 2021), promising to extend findings and theory from classical neuroanatomy in non-human animals to humans. Yet, as most tools generally focus on the processing of individual modalities, or the combination of at most two different modalities (e.g. T1-weighted MRI and rs-fMRI), researchers interested in additional synergies across an even larger catalogue of modalities are forced to develop custom-built image co-registration and data-integration procedures.

System neuroscience has increasingly benefitted from paradigms that combine different imaging modalities (Paquola et al., 2020; Van den Heuvel et al., 2019; Van den Heuvel and Yeo, 2017). For example, multiple studies have begun to study brain function and functional connectivity in surface-based anatomical reference frames (Huntenburg et al., 2021; Tierney et al., 2013; Vos de Wael et al., 2018), and combined these assessments with diffusion MRI approaches (Liu et al., 2016; Hong et al., 2019). Further work integrating structural and functional neuroimaging modalities has propelled interest in examining structure-function relationships in the human brain (Huntenburg et al., 2018; Suárez et al., 2020; Benkarim et al., 2021; Paquola et al., 2019b; Vázquez-Rodríguez et al., 2019). Furthermore, there has been significant development towards the identification of multimodal parcellations (Fan et al., 2016; Eickhoff et al., 2018; Genon et al., 2021, 2018; Glasser et al., 2016) and large-scale gradients of brain organization (Vos de Wael et al., 2020, 2021; Margulies et al., 2016; Paquola et al., 2020, 2019a, 2019b; Valk et al., 2020; Müller et al., 2020; Tian et al., 2020).

To build upon existing MRI processing pipelines that are primarily geared towards single modalities, we developed *micapipe* (http://micapipe.readthedocs.io). The pipeline integrates advanced processing streams for structural MRI, resting-state functional MRI (rs-fMRI), diffusion-weighted MRI, and myelin-sensitive MRI to automatically generate models of structural, functional, and microstructural human brain organization. *Micapipe* generates inter-regional matrices across different spatial scales, using several cortical as well as subcortical parcellations (Desikan et al., 2006; Destrieux et al., 2010; Scholtens et al., 2018; von Economo, 2009; Fischl, 2012; Vos de Wael et al., 2020; Schäfer et al., 2018; Glasser et al., 2016; Patenaude et al., 2011; Diedrichsen et al., 2009). In a nutshell, *micapipe* transforms BIDS-conform MRI data (Gorgolewski et al., 2017) to processed macroscale connectomes in an easy-to-analyze format. Easy-to-verify outputs and visualizations can be produced for quality control (QC). In addition to its codebase being openly available on GitHub (http://github.com/MICA-MNI/micapipe), *micapipe* is also available as a container (Docker, included as BIDS App), and is accompanied by detailed tutorials and documentation.

## 2. Results

*Micapipe* has a modular workflow that can incorporate multiple MRI data modalities (T1-weighted MRI, myelin-sensitive MRI, diffusion-weighted MRI, and resting-state functional MRI), converting BIDS-conform input into BIDS-conform surface, volume, and matrix data (**Figure 1A**). The following sections describe key pipeline features, main outputs, and detail automated quality control (QC) visualizations. We also perform several validation experiments across a diverse range of datasets.

**Figure 1.**
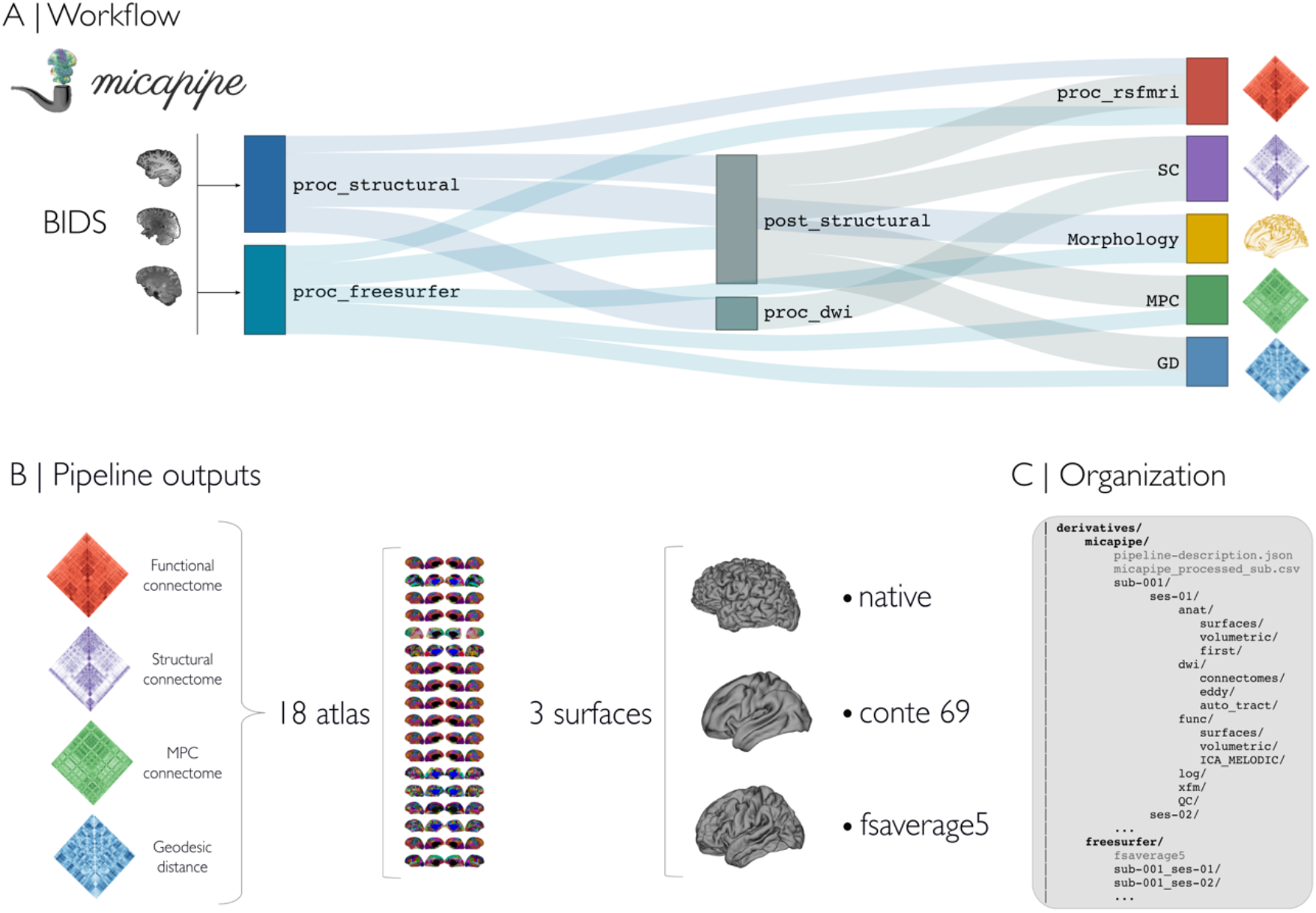
**A)** Pipeline workflow. **B)** Outputs can be generated across 18 different cortical parcellations (100-1000 parcels), in addition to subcortical and cerebellar parcellations. Most results are mapped to three different surface spaces: native, conte69 and fsaverage5. **C)** Outputs are hierarchically ordered with BIDS-conform naming.

### 2.1 Pipeline workflow

Processing modules of *micapipe* can be run individually or bundled using specific flags via a commandline interface. Multimodal integration relies strongly on characterization of anatomy via the processing of T1-weighted MRI data. Using volume and surface-based processing streams, subcortical, cortical and cerebellar segmentations are generated in subject- and modality-specific spaces. Using structural imaging data, in addition to other input modalities, inter-regional brain matrices can be generated across 18 combinations of cortical, subcortical, and cerebellar parcellations. Inter-regional matrices are: i) structural connectomes (SC) derived from diffusion tractography (Smith et al., 2015a), ii) functional connectomes (FC) derived from resting-state signal correlations (Biswal et al., 2010), iii) geodesic distance (GD) matrices that quantify cortico-cortical proximity using cortical surface models (Ecker et al., 2013; Hong et al., 2018), and iv) microstructural profile covariance (MPC) matrices that assess inter-regional similarity in intracortical intensity profiles from microstructurally-sensitive imaging (Paquola et al., 2019b). Surface-mapped features are made available across three surfaces (**Figure 1B**): native, conte69 (Van Essen et al., 2012), and fsaverage5 (Fischl et al., 1999). Intermediary files and processed derivatives and matrices conform to BIDS naming conventions (**Figure 1C**), facilitating future use and harmonization across datasets and software.

### 2.2 Quality control (QC)

The QC module visualizes outputs at the individual and group levels (**Figure 2A**). Reports detail completed processing steps, including image registrations, surface parcellations, and region-to-region matrices. They are organized by modality and parcellation. These reports help users to identify missing data, poor image quality, and faulty registrations (**Figure 2A**). Complementing subject-specific reports, group level QC automatically generates a report outlining completed and missing modules for each subject facilitating use for large datasets (**Figure 2B**).

**Figure 2.**
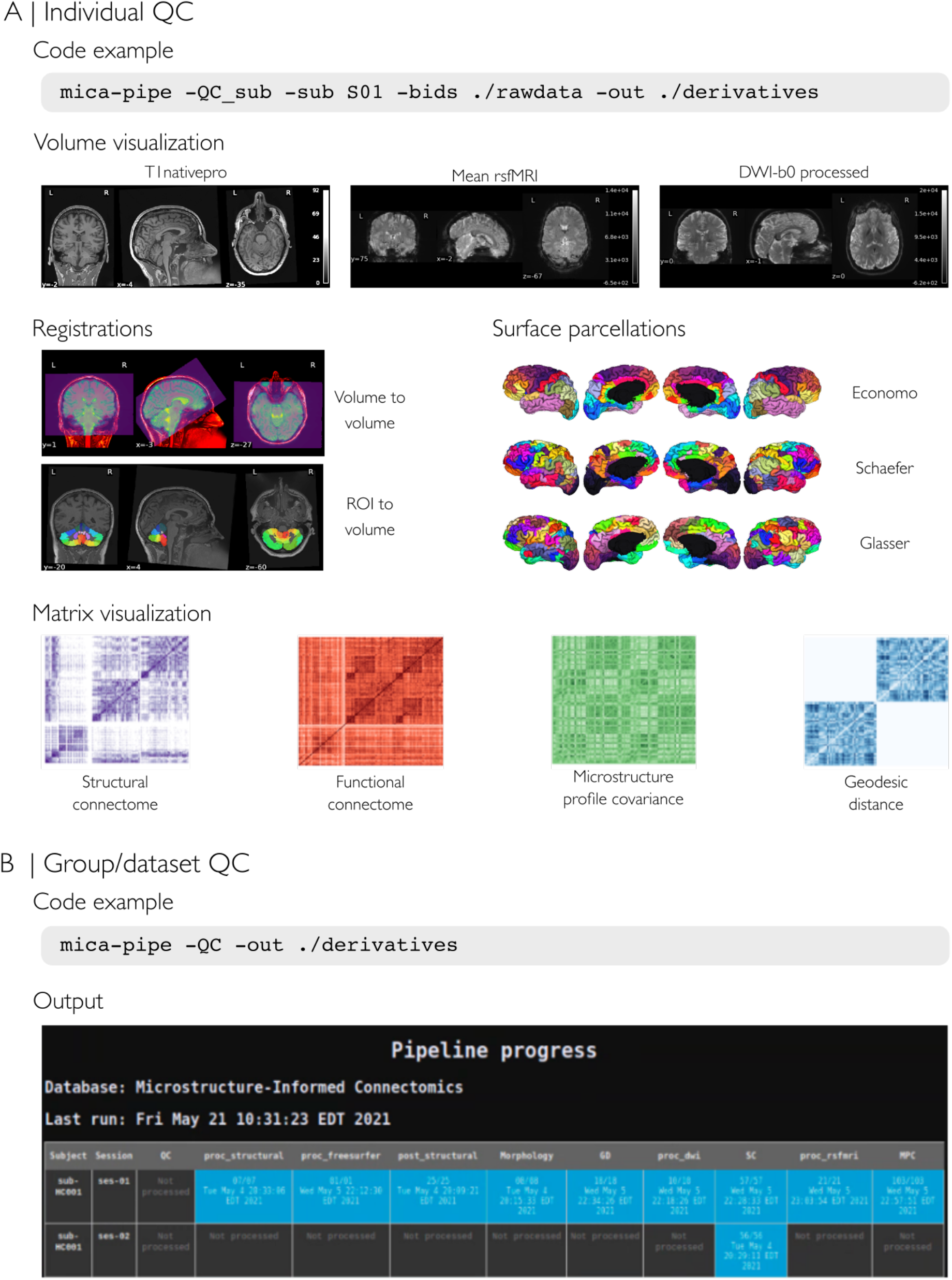
**A)** Individual level quality control (QC), which can be run at any point during the processing. The QC procedure will generate a html report file for each subject containing visualizations of intermediate files for volume visualization, cross-modal co-registrations, and surface parcellations. Moreover, it allows inspection of inter-regional matrices such as the structural connectome (from diffusion MRI tractography), the functional connectome (from resting-state fMRI signal correlation), the microstructural profile covariance matrix (from correlations of intracortical microstructural profiles), and geodesic distance matrices. **B)** QC can also be run at a group/dataset-level. The report consists of a color-coded table with rows as subjects and columns as the pipeline modules (*blue*: completed, *orange*: incomplete/error, *dark gray*: not processed).

### 2.3 Assessing output consistency within and between datasets

We evaluated whether *micapipe* yields consistent results across 50 individuals of an openly available multimodal MRI dataset [MICA-MICs; (Royer et al., 2021), and also compared processed outputs to those from six additional datasets (**Table S1**).

We first assessed within-dataset consistency for each modality (GD, SC, FC, MPC) at three different granularities (Schäfer 100, 400 and 1000 parcels) using five different metrics. We generated modality- and dataset-specific mean group matrices and computed consistency across the following features: the first eigenvector/gradient explaining the most data variance (calculated via diffusion map embedding (Coifman et al., 2006)), the matrix edges, as well as node strength, characteristic path length, and clustering coefficient as three representative graph features (Rubinov and Sporns, 2010), **Figure 3A**]. We correlated subject-level and group-level metrics to quantify within-dataset consistency (Spearman’s rho, see **Supplementary Figure 3A**). Correlations were highest for GD and SC, followed by FC and MPC. Gradient 1 was the most consistent measure across parcellations and modalities, followed by edges and node strength. Overall, characteristic path length and clustering coefficient were similar at lower granularity (100 parcels) but increasingly dissimilar at higher granularity (1000 parcels). Findings were consistently observed across all datasets (**Figure 3B-C**).

**Figure 3.**
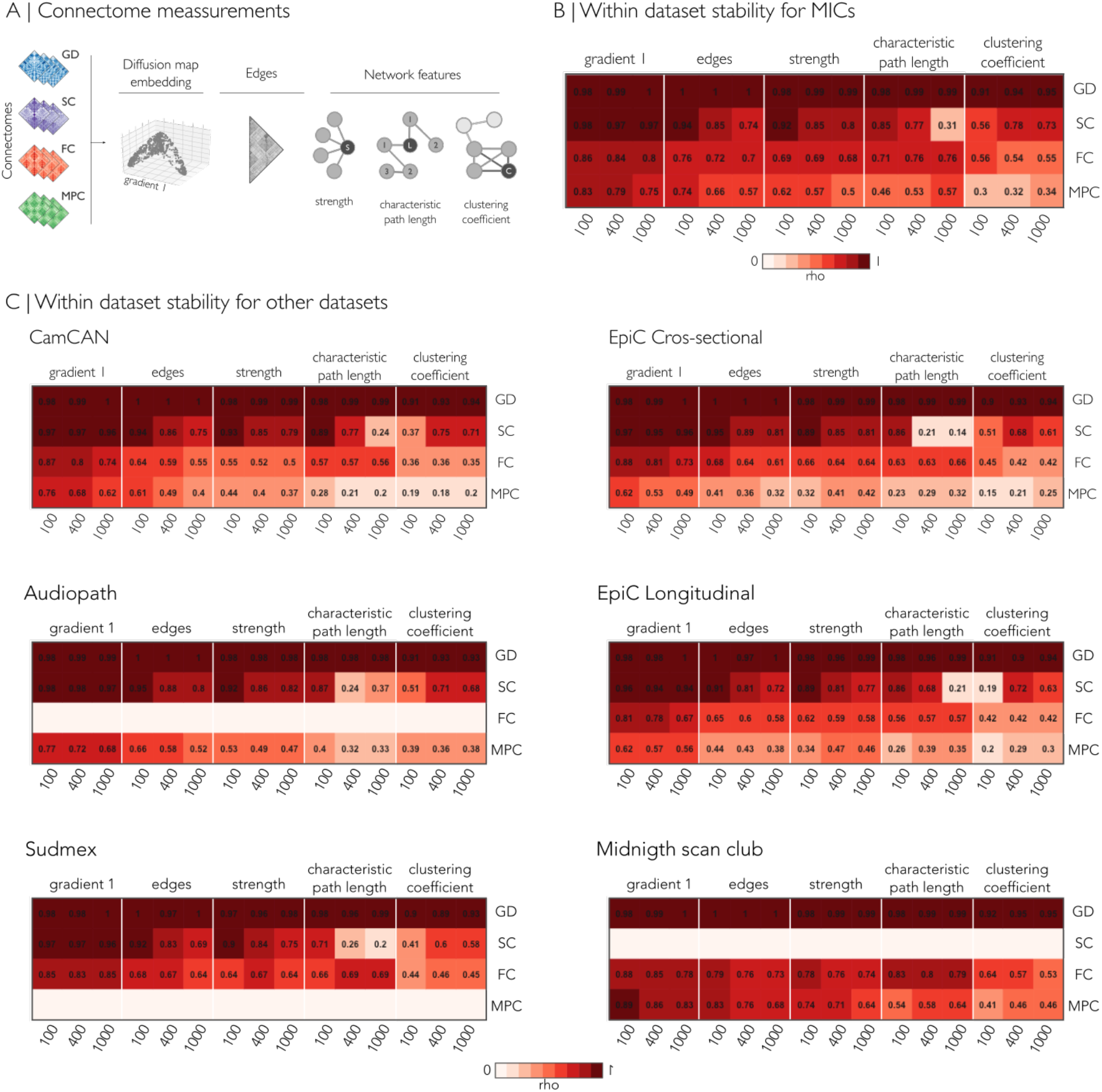
Mean consistency value, indicating the Spearman’s rho between subject- and the group-level measurements, for the Schäfer-100, Schäfer-400 and Schäfer-1000 parcellations. **A)** For each modality, five measurements were evaluated: principal gradient, edges, node strength, path length, and clustering. Empty rows indicate modalities that were not analyzed. MPC: microstructural profile covariance, FC: functional connectivity, SC structural connectivity, GD geodesic distance

We also compared between datasets (**Figure 4**). As for the within-dataset analysis, we found the highest consistency between datasets for GD and SC, followed by FC and MPC. GD, SC and FC showed high similarity between datasets for the edges, first eigenvector/gradient, and node strength. FC had decreased consistency between datasets for characteristic path length and clustering coefficient. MPC had the lowest between dataset consistency for all measurements.

**Figure 4.**
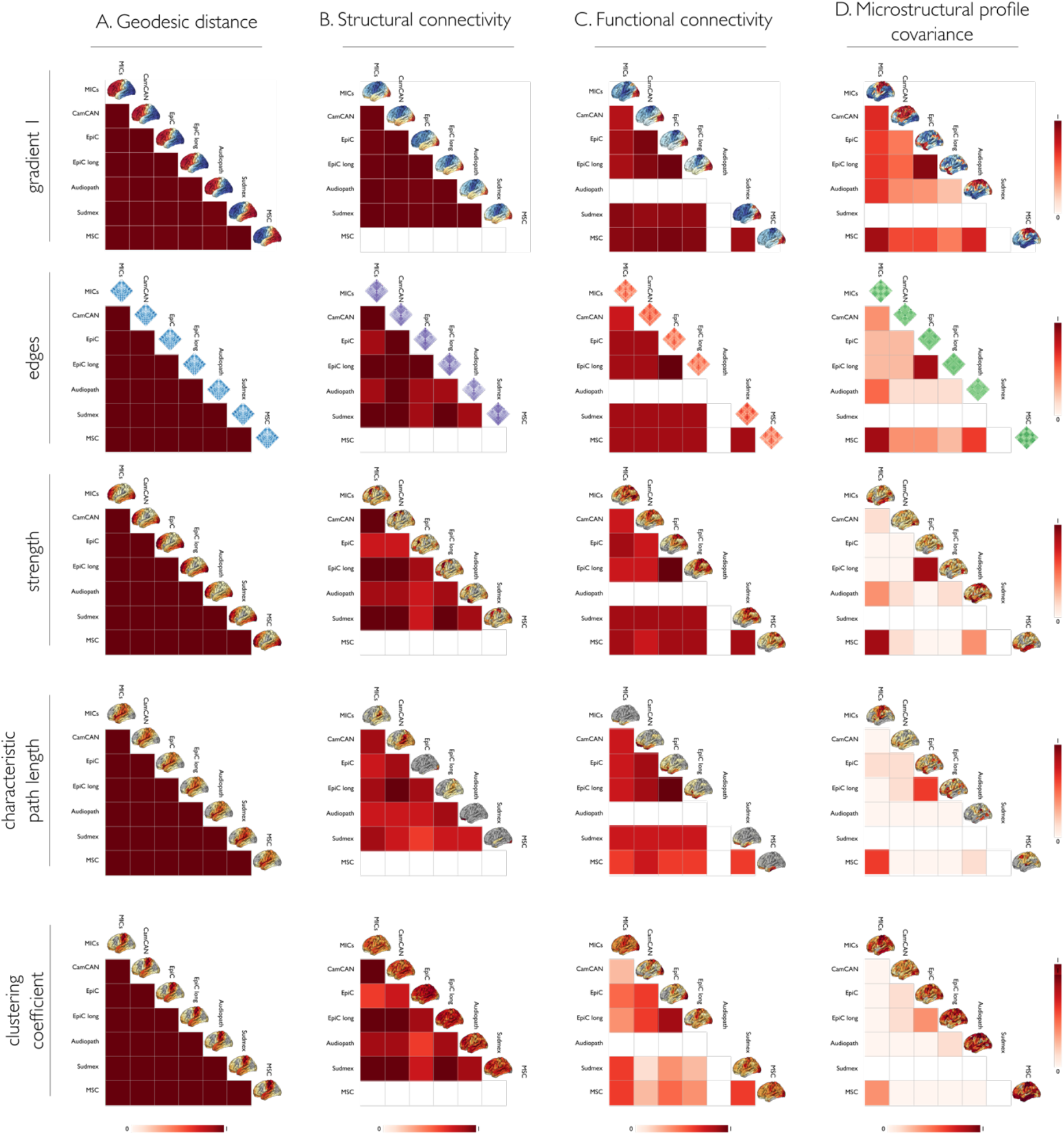
We assessed consistency of matrix parameters across datasets using Spearman’s rho correlation coefficient for the same features as in Figure 3. Each column represents the different modality connectomes: A) geodesic distance, B) structural connectome, C) functional connectome, and D) microstructural profile covariance.

### 2.4 Performance

We tested *micapipe* on seven different databases acquired using different MRI sequence/parameter combinations (**Table S1**). Processing times varied depending on the image resolution, the need to additionally process data using FreeSurfer, the number of streamlines selected for the structural connectome generation, and the type of acquisitions per dataset (**Table S2**). Processing was performed on the Brain Imaging Center (BIC) cluster of the Montreal Neurological Institute and Hospital on Ubuntu 18.04.5 LTS version workstations. A maximum virtual memory of 6GB, with 6-10 CPU cores, and 20 GB of RAM were required. Output size depended on image resolution and the length of the rs-fMRI acquisitions (**Table S3**).

### 2.5 Software and data availability

An expandable documentation at https://micapipe.readthedocs.io describes installation, usage, pipeline steps, updates, extra features, and provides a series of ready-to-use tutorials. All code can be found at https://github.com/MICA-MNI/micapipe, and is published under the General Public License 3.0. *Micapipe* is delivered as a docker container via BIDS-App [http://bids-apps.neuroimaging.io/apps/ (Gorgolewski et al., 2017)], and available on ReproNim [https://github.com/ReproNim/containers (Halchenko et al., 2021)]. Detailed steps to use the Docker container and to build a corresponding singularity container are available under the readthedocs documentation. Code for figures and tables can be found in the *micapipe-supplementary* GitHub repository (https://github.com/MICA-MNI/micapipe-supplementary).

## 3. Discussion

We present *micapipe*, an open software package to integrate and process raw multimodal MRI data into a range of multiple measures of structural and functional human brain network organization. As a standalone and BIDS App, *micapipe* inputs and outputs BIDS-conform MRI data. Its outputs consist of derivative features across multiple parcellations, available in both surface- and volume-based reference spaces. Notably, *micapipe* outputs include measures of brain morphology, together with inter-regional matrices encoding cortico-cortical spatial proximity (based on geodesic distance analysis along cortical surfaces derived from T1-weighted MRI), intrinsic functional connectivity (based on rs-fMRI signal correlations), structural connectivity (derived from diffusion MRI tractography), and microstructural similarity (derived from intracortical profile covariance analysis of myelin-sensitive MRI) aggregated across 18 different cortical as well as subcortical/cerebellar parcellations. Given the complexity of multimodal processing and analysis, *micapipe* furthermore offers advanced functionalities for individual and group-level QC at different stages of processing as well as final outputs. As a unified tool to fuse and analyze multimodal neuroimaging data, *micapipe* offers neuroscientists a workflow to robustly interrogate human brain organization across multiple scales.

A range of pipelines have previously been developed to process MRI images from specific modalities, including tools for the generation of cortical and subcortical segmentations based on T1-weighted MRI data (Das et al., 2009; Fischl, 2012; Kim et al., 2005), pipelines to process functional MRI data (Craddock et al., 2013; Esteban et al., 2019), as well as tools for diffusion MRI data handling (Cieslak et al., 2021; Jenkinson et al., 2012; Tournier et al., 2019). Several workflows have furthermore been developed for connectome mapping (Daducci et al., 2012; Whitfield-Gabrieli and Nieto-Castanon, 2012), which allow users to examine structural and functional network architecture in a systematic manner. Building upon these developments, *micapipe* offers a unified framework for multimodal fusion and data processing. As such, it is similar in scope to the proposed Connectome Mapper tool (Daducci et al., 2012), although with notable differences. In particular, *micapipe* incorporates a stream for the surface-based mapping of intracortical myelin proxies and for the generation of microstructural profile covariance (Paquola et al., 2019b). A growing body of literature emphasizes the utility of myelin-sensitive MRI analysis for cortical parcellation (Carey et al., 2018; Glasser and Van Essen, 2011; Granberg et al., 2017), to assess brain and cognitive development (Deoni et al., 2012; Whitaker et al., 2016; Lebel and Deoni, 2018; Paquola et al., 2019a), and to interrogate microstructural imbalances in common brain disorders (Cooper et al., 2019; Du et al., 2019; Bernhardt et al., 2018; Larivière et al., 2019). Recent work has shown that the analysis of covariance patterns of intracortical microstructural profiles can generate new descriptions of large-scale network organization (Paquola et al., 2020; Royer et al., 2020). These networks appear to be primarily governed by systematic shifts in laminar differentiation and neuronal density, showing a principal organizational axes similar to those at the level of cytoarchitecture and intrinsic functional connectivity (Margulies et al., 2016; Paquola et al., 2021, 2019b). A further notable feature is the automated generation of cortico-cortical geodesic distance matrices, which indexes proximity between different regions on the folded cortical surface. Cortico-cortical geodesic distance has been suggested to relate to intrinsic, horizontal connectivity within the cortical ribbon as well as to cortical wiring cost (Ecker et al., 2013; Hong et al., 2018; Paquola et al., 2020). Moreover, several investigations into principles of macroscale brain organization have emphasized that the brain is a physically embedded network, and that thus inter-regional distance relationships may help in the understanding of the topographic layout of functional systems and the connections formed between them (Betzel et al., 2016; Betzel and Bassett, 2018; Margulies et al., 2016; Valk et al., 2020; Wang et al., 2021; Smallwood et al., 2021).

A series of evaluations assessed consistency of *micapipe* outputs, studying data from 455 individuals across seven datasets. Our evaluations focused on the consistency of inter-regional matrix edges, the first eigenvector (gradient), and three widely used graph theoretical measurements (node strength, characteristic path length, clustering coefficient) for up to four matrix modalities (i.e., geodesic distance, functional connectivity, structural connectivity, microstructural covariance). Altogether, our results indicate a generally high consistency of the first gradient across datasets, with some variations across modalities. For example, geodesic distance and structural connectivity gradients were markedly consistent (r>0.95), followed by functional connectivity and microstructural profile covariance. It is likely that functional connectivity measures and associated gradients may, in part, be influenced by state-to-state variations compared to the more static measures of structural connectivity and geodesic distance, likely in addition to data acquisition related effects. Edges and graph derived measurements followed an analogous pattern of consistency. With respect to the relatively low stability of microstructural profile covariance, one needs to highlight that the included datasets greatly varied in terms of microstructurally sensitive MRI contrasts, featuring T1-weighted/T2w intensity ratio (Glasser and Van Essen, 2011), quantitative T1 relaxometry (Royer et al., 2021), as well as magnetization transfer imaging (Shafto et al., 2014). While these sequences are all though to be sensitive to intracortical myelin content, their individual biophysical specificity remains to be established.

Through the successful integration of several processing tools, *micapipe* provides multiple ready to use inter-regional feature matrices, i.e., structural connectomes, functional connectomes, microstructural covariance, and geodesic distance, together with QC procedures. Our pipeline is supported by a growing ecosystem of open tools for data and code sharing, notably Github, readthedocs, Docker, BIDS Apps (Gorgolewski et al., 2017), and repronim/datalad (Robert et al., 2016). By making *micapipe* openly accessible as well, we hope that it will be beneficial for future studies on human brain organization.

## 4. MATERIALS AND METHODS

*Micapipe* runs modular processing streams on BIDS-conform raw *T1-weighted*, *microstructure-sensitive*, *diffusion-weighted*, and *resting-state functional MRI* data to generate fully processed surface/volume features as well as inter-regional feature matrices. A documentation with detailed descriptions on the installation, implementation, as well as usage examples and output files are available at https://micapipe.readthedocs.io/.

### 4.1 Workflow and main processing modules

*Micapipe* requires the input dataset to be formatted in BIDS (Gorgolewski et al., 2016).

#### Structural processing

Structural processing operates on T1-weighted images. The structural processing workflow can perform volumetric (with command-line option: -*proc_structural*) and surface-based *(-proc_freesurfer, - post_structural, -GD, -Morphology*) processing. The workflow registers subject data to volumetric and surface-templates providing several useful structural metrics for further analyses. These include geodesic distance matrices (-*GD*) mapped to multiple parcellation schemes as well as vertex-wise cortical thickness and curvature data (-*Morphology*). The structural workflow includes tools from AFNI (Cox, 1996), FSL (Jenkinson et al., 2012), ANTs (Avants et al., 2011), Mrtrix3 (Tournier et al., 2019) and FreeSurfer (Fischl, 2012). Further information about the usage and outputs is found in the structural processing section in the online documentation.

##### Proc_structural

Initial structural pre-processing (i.e., *-proc_structural)* keeps all data in volumetric format and generates a T1-weighted image in native processing space (nativepro, **Figure S1A**). Each T1-weighted run is reoriented to LPI orientation (*i.e*., left-right, posterior-anterior, inferior-superior), de-obliqued, and oriented to standard space (MNI152). If multiple T1-weighted scans are found in the raw data, they are linearly aligned to the first run and averaged. Next, the average image is corrected for intensity nonuniformity (N4, Tustison et al., 2010) and intensity is normalized between 0 and 100. The resulting image is named T1nativepro, which stands for ***T1***-weighted in ***native pro***cessing space. ***T1nativepro*** is skull-stripped, subcortical structures are segmented using FSL FIRST (Patenaude et al., 2011), and tissue types are classified (gray matter, white matter, CSF) using FSL FAST (Zhang et al., 2001). A non-linear registration to MNI152 (0.8mm and 2mm resolutions) is calculated (Tustison and Avants, 2013) and a five-tissue-type (5TT) image segmentation is generated for anatomically constrained tractography.

##### Proc_freesurfer

Cortical surface segmentations are generated from native T1-weighted scans using FreeSurfer 6.0 (Fischl, 2012) and ordered under the FreeSurfer directory, following BIDS naming conventions. We provide an option for datasets that have already been quality controlled to easily integrate the results within the pipeline’s directory structure and an option to process with voxel sizes less than 1mm^3^ at native resolution (-*hires*). We recommend that users carefully inspect and, if needed, manually correct FreeSurfer-generated cortical surface segmentations. As *micapipe* relies heavily on surface-based processing, poor segmentation quality may compromise downstream results.

##### Post_structural

The first step of the post structural processing is to calculate an affine registration from native FreeSurfer space to *T1nativepro* space. It then registers a probabilistic cerebellar atlas (Diedrichsen et al., 2009) from MNI152 to the subject’s T1nativepro space using affine and non-linear transformations previously computed in the *-proc_structural* module. Next, a surface-based registration of fsaverage5 annotation labels to native surface is performed, and the surface-based parcellation in native FreeSurfer space is transformed into a volume. Finally, the transformation matrices are applied to bring each volumetric parcellation from native FreeSurfer space to *T1nativepro* space. In the last step of this module, the pipeline builds a conte69-32k sphere and resamples white, pial and midthickness native surfaces to the conte69-32k template.

The *-post_structural* module registers native FreeSurfer-space cortical surfaces to two different standard templates (fsaverage5 and conte69), in addition to mapping all cortical parcellation schemes to the subject’s native surface space and volumetric *T1nativepro* space (**Figure S1B**). *Micapipe* provides a total of 18 cortical, subcortical and cerebellar parcellations at different resolutions according to anatomical, cytoarchitectural, intrinsic functional, and multimodal schemes, at different resolutions. Anatomical atlases available in *micapipe* include Desikan-Killiany (aparc, Desikan et al., 2006) and Destrieux (aparc.a2009s, Destrieux et al., 2010) parcellations provided by FreeSurfer, as well as an *in vivo* approximation of the cytoarchitectonic parcellation studies of Von Economo and Koskinas (Scholtens et al., 2018). Additionally, we include similarly sized sub-parcellations, constrained within the boundaries of the Desikan-Killiany atlas, providing matrices with 100, 200, 300, and 400 cortical parcels following major sulco-gyral landmarks (Fischl, 2012; Vos de Wael et al., 2020). Parcellations based on intrinsic functional activity are also included across several granularities (100, 200, 300, 400, 500, 600, 700, 800, 900, and 1000 nodes, (Schäfer et al., 2018). Lastly, we also provide a multimodal atlas with 360 nodes derived from the Human Connectome Project dataset (Glasser et al., 2016). All atlases are provided on Conte69 and fsaverage5 surface templates, and on each participant’s native surface to generate modality-specific matrices in subsequent modules.

##### Morphology

This module registers cortical thickness and curvature measurements to two distinct templates. Both surface-based morphological features are registered to fsaverage5 and conte69 and smoothed with a gaussian filter with full width half maximum of 10mm.

##### GD: geodesic distance

Individual GD matrices are computed along each participant’s native cortical midsurface using workbench tools (Marcus et al., 2011). First, a centroid vertex is defined for each cortical parcel by identifying the vertex with the shortest summed Euclidean distance from all other vertices within its assigned parcel. Then, the geodesic distance is calculated from the centroid vertex to all other vertices on the midthickness mesh using Dijkstra’s algorithm (Dijkstra, 1959). Notably, this implementation computes distances not only across vertices sharing a direct connection, but also across pairs of triangles which share an edge in order, thus mitigating the impact of mesh configuration on calculated distances. Vertex-wise GD values were averaged within parcels to improve computation performance.

#### Diffusion-weighted imaging processing

This section describes all DWI-related processing steps implemented in *micapipe*, which heavily rely on tools from MRtrix3 (Tournier et al., 2019). This includes image processing preparation for the construction of tractography-based structural connectivity matrices, as well as associated edge length matrices, all in native DWI space (**Figure S1C)**. *Micapipe* DWI processing has been optimized for multishell DWI but can also handle single-shell data. Geometric and inhomogeneity corrections are performed in datasets that contain one or more reverse phase encoding DWI. It is a mandatory requirement that all DWIs have associated bvec, bval and json files, with encoded phase direction and total readout time.

##### Proc_dwi

This module processes DWI scans, and derives several potentially useful metrics (e.g., fractional anisotropy, mean diffusivity, **Figure S1C**). First, if there is more than one set of DWI scans in the BIDS directory, they are aligned to each other using a rigid-body registration, and concatenated. All DWI images are then converted to Mrtrix Imaging format (mif), which encodes the bvec, bval, phase encoding direction and total readout time. Concatenated DWI images undergo denoising by estimating data redundancy in the PCA domain via a Marchenko-Pasteur approach [MP-PCA, (Veraart et al., 2016; Cordero-Grande et al., 2019)]. Then, Gibbs ringing artifact correction is applied (Kellner et al., 2016), and residuals are calculated from denoised images for QC purposes. Provided a reverse phase encoding, susceptibility distortion, head motion, and eddy currents are corrected (Andersson et al., 2003; Smith et al., 2004; Andersson and Sotiropoulos, 2016). If none is provided, only motion correction is performed. Additionally, outlier detection and replacement are applied (Andersson et al., 2016). After this step, the quality of the motion and inhomogeneity corrected diffusion images is assessed using eddy_quad (Bastiani et al., 2019), and a non-uniformity bias field correction (Tustison et al., 2010) is applied to finalize DWI preprocessing. Next, the b0 image is extracted from the corrected DWI and linearly registered to the main structural image (i.e., *T1nativepro*). A DWI brain mask is generated by registering the MNI152 brain mask to DWI space using previously generated transformations. A diffusion tensor model (Basser et al., 1994) is then applied to the corrected DWI and the fractional anisotropy and mean diffusivity images are computed (Veraart et al., 2013). An estimation of the response function is calculated over different tissues: cerebrospinal fluid, white matter, and gray matter (Dhollander, et al., 2016). These are later used to estimate the fiber orientation distribution (FOD) by spherical deconvolution (Jeurissen et al., 2014; Tournier et al., 2004). Next, intensity normalization is applied to each tissue FOD (Raffelt, et al., 2017). A second registration, in this case non-linear, is calculated between the normalized white matter FOD and the T1-weighted image previously registered linearly to DWI space. The resulting warp field allows for an improved registration between the T1-weighted and the native DWI space in most datasets. Finally, the 5TT segmentation image is registered to native DWI space and a gray matter white matter interface mask is calculated. For QC purposes, a track density image (Calamante et al., 2010) is computed with 1 million streamlines using the iFOD1 algorithm (Tournier et al., 2012) and anatomically constrained tractography (Smith et al., 2012).

##### SC: structural connectome generation

Structural connectomes are generated with Mrtrix3 from pre-processed DWI data from the previous module and subcortical and cerebellar parcellations are non-linearly registered to native DWI space. First, a tractography with 40 million streamlines (default but modifiable, maximum tract length=400, minimum length=10, cutoff=0.06, step=0.5) is generated using the iFOD2 algorithm and 3-tissue anatomically constrained tractography (Smith et al., 2012; Tournier et al., 2010). A second tract density image (TDI) of the resulting tractography is computed for QC. By default, the full brain tractography is erased at the end of this module but can be kept using the option “-*keep_tck*”. Next, spherical deconvolution informed filtering of tractograms [SIFT2 (Smith et al., 2015a)] is applied to reconstruct whole brain streamlines weighted by cross-sectional multipliers. The reconstructed cross-section weighted streamlines are then mapped to each parcellation scheme, with (i) cortical, (ii) cortical and subcortical, and (iii) cortical, subcortical, and cerebellar regions (Smith et al., 2015b. These are also warped to DWI native space. The connection weights between nodes are defined as the weighted streamline count, and edge length matrices are also generated.

#### Resting-state fMRI processing

This module processes the rs-fMRI scans, in preparation for the construction of functional connectomes. This pipeline is optimized for spin-echo images with reverse phase encoding used for distortion correction. The pipeline is mainly based on tools from FSL and AFNI for volumetric processing, as well as FreeSurfer and Workbench for surface-based mapping (**Figure S1D)**.

Initial fMRI processing steps involve the removal of the first five volumes to ensure magnetic field saturation, image re-orientation (LPI), as well as motion and distortion correction. Motion correction is performed by registering all time-point volumes to the mean volume, while distortion correction leverages main phase and reverse phase field maps acquired alongside rs-fMRI scans. Nuisance variable signal is removed either using an ICA-FIX classifier with a default training set or custom training set input by the user (Griffanti et al., 2014; Salimi-Khorshidi et al., 2014) or by selecting white matter, CSF, and global signal regression [for Discussion, see (Murphy et al, 2009, Murphy & Fox 2017, Vos de Wael et al. 2017)]. Additionally, a regression of time points with motion spikes is performed using motion outlier outputs provided by FSL. Volumetric timeseries are averaged for registration to native FreeSurfer space using boundary-based registration (Greve and Fischl, 2009), and mapped to individual surface models using trilinear interpolation. Native-surface and template-mapped cortical time series undergo spatial smoothing (Gaussian kernel, FWHM = 10mm), and are subsequently averaged within nodes defined by several parcellation schemes. Parcellated subcortical and cerebellar time series are also provided and are appended before cortical time series.

##### FC: functional connectome generation

Individual rs-fMRI time series are mapped to individual surface models. Native surface-mapped time series are registered to standard surface templates (fsaverage5, conte69). Native surface and conte69-mapped time series are averaged within cortical parcels. The subcortical and cerebellar parcellations are warped to each subject’s native rs-fMRI volume space and used to extract the time series within each node. Individual functional connectomes are generated by cross correlating all nodal time series.

#### Microstructural processing and microstructural profile covariance (MPC) matrix generation

This module samples intracortical intensities from a quantitative MRI contrast, generating a depth-dependent intracortical intensity profile at each vertex of the native surface mesh. By parcellating and cross-correlating nodal intensities, this module generates MPC matrices. This approach has been previously applied over the whole cortex (Paquola et al., 2019b), as well as in targeted structures such as the insula (Royer et al., 2020). The first processing step is a boundary-based registration from the quantitative imaging volume (default) or input microstructurally sensitive image contrast to FreeSurfer native space. Then, 16 equivolumetric surfaces (Waehnert et al., 2014) are generated between the pial, and white matter boundary previously defined from FreeSurfer. Intracortical equivolumetric surfaces are generated using https://github.com/kwagstyl/surface_tools. Surfaces closest to pial and white matter boundaries are discarded to minimize partial volume effects, resulting in 14 surfaces. A surface-based registration is performed to fsaverage5 and conte69-32k templates, and the vertex-wise intensity profiles are averaged within parcels for each parcellation. Nodal profiles are cross-correlated across the cortical mantle using partial correlations controlling for the average cortex-wide intensity profile. Several regions are excluded when averaging cortex-wide intensity profiles, including left/right medial walls, as well as non-cortical areas such as the corpus callosum and pericallosal regions.

### 4.2 Quality Control

*Micapipe* includes an integrated QC module, which can be run at any point during processing. This step generates group-level and individual-specific QC reports allowing the user to identify missing files, verify registration performance, and check outputs requiring further inspection. Individual QC generates an html report with detailed information of each processing step (**Figure 2A**). The report contains different tabs, one per module: main inputs and outputs of each module, main parameters of the processing steps (obtained from the metadata json sidecar files generated by *micapipe*), volume visualization of the main outputs, visualization of the main registrations, different surfaces generated by the pipeline, parcellations plotted on native surface, structural connectome matrices, functional connectome matrices, geodesic distance matrices, microstructural intensity profiles and connectomes and microstructural profiles (image intensities at each cortical depth) plotted on the native surface. Group level QC generates a color-coded table (with rows for subjects and columns for modules (**Figure 2B**)).

### 4.3 Additional features

#### Automatic bundle segmentation

The *micapipe* repository also includes an optional *<u>automatic virtual dissection</u>* of major fiber tracts (**Figure S2A**). This tool is an adaptation of XTRACT (Warrington et al., 2020) implemented using Mrtrix3 and ANTs, and its main purpose is to split a tractography (tck file) into the main white matter tracts. The automatic bundle segmentation uses already established automatic dissection protocols manually tuned for optimal performance. Derived from a full brain tractography, 35 bundles are virtually dissected using the *<u>LANIREM protocols</u>*. The quality of the full brain tractography will determine the quality of bundle separation. It is highly recommended to provide a tractography with more than one million streamlines, and QC for any errors. Strategies such as anatomically constrained tractography (ACT) and spherical deconvolution informed filtering of tractographies (SIFT), which are available in Mrtrix3, should aid in obtaining such high-quality tractographies. For processing a Nonlinear (SyN) registration of the native FA map to the FA atlas (FMRIB58_FA_1mm) is calculated. Resulting transformations are then applied to each bundle protocol to register them to the native FA space (DWI). Finally, each white matter bundle is filtered according to the dissection protocols.

#### Anonymize function

A function to anonymize the anatomical images from the BIDS directory for data sharing is provided within the *micapipe* repository as an extra feature. Native structural images are anonymized and deidentified with one of three different methods: de-facing, linear refacing or refacing with a non-linear warp field (**Figure S2B**). This tool uses a custom template and a set of ROIs specifically developed to identify the face and skull. The full head template was created using the T1-weighted images (resolution of 0.8×0.8×0.8mm) of 60 randomly selected healthy individuals from the MICA-MICs dataset (Royer et al., 2021). An inter-subject non-linear registration was performed without any mask, then the template was built using the mean of the normalized images. Three masks were generated: an ROI that covers the face, a brain mask, and a brain and neck mask. Unlike other algorithms, *micapipe_anonymize* supports different anatomical modalities (*e.g*., quantitative T1 maps).

### 4.4 Feature matrices

Besides surfaces and parcellations, *micapipe* outputs up to four inter-regional matrices across several parcellation: structural connectome (SC), functional connectome (FC), geodesic distance (GD) and microstructural profile covariance (MPC). Rows and columns of GD and MPC matrices follow the order defined by annotation labels associated with their parcellation (see *<u>parcellations</u>* in the *micapipe* repository), including unique entries for the left and right medial walls. For example, row and column entries of the Schäfer-100 matrices are ordered according to: Left hemisphere cortical parcels (1 medial wall followed by 50 cortical regions), and right hemisphere cortical parcels (1 medial wall followed by 50 cortical regions). FC and SC matrices follow the same ordering, although entries for subcortical and cerebellar structures are appended before cortical parcels. As such, row and column entries of the Schäfer-100 FC and SC matrices are ordered according to: Subcortical structures and hippocampus (7 left, 7 right), cerebellar nodes (34 regions), left hemisphere cortical parcels (1 medial wall followed by 50 cortical regions), and right hemisphere cortical parcels (1 medial wall followed by 50 cortical regions). The ordering of all parcels and their corresponding label in each volumetric parcellation are documented in lookup tables provided in the *micapipe* repository (*<u>parcellations/lut</u>*). Further information about the organization and visualization of the output connectomes can be found in the respective section of the *<u>documentation</u>*.

### 4.5 Validation experiments

The pipeline was tested in 455 human participants from seven datasets (**Table S1, S4**): MICA-MICs, (Royer et al., 2021), EpiC-UNAM (Rodríguez-Cruces et al., 2020), Cam-CAN (Shafto et al., 2014), SUDMEX (Angeles-Valdez et al., 2021), MSC (Gordon et al., 2017), and 7T-Audiopath (Sitek et al., 2019). EpiC-UNAM consists of two separate acquisitions: one cross-sectional and one longitudinal. Acquisition and processing details for each dataset can be found in the section “Processing databases” of the online documentation.

#### Inter-subject consistency

We assessed inter-subject consistency at the level of the first eigenvector/gradient of each matrix, matrix edges, and three widely graph theoretical measures (node strength, characteristic path length, clustering coefficient). Evaluations were carried out across three selected parcellations(Schäfer-100, 400 and 1000). Inter-subject consistency was quantified as the Spearman correlation between each participant measure and the group average measure for each available modality. This procedure was applied for the gradient 1, edges, and the three graph features (**Figure 3**).

To generate *gradients*, we used BrainSpace (http://brainspace.readthedocs.io, Vos de Wael et al., 2020), with the following options: normalized angle kernel, diffusion embedding with alpha=0.5 and automatic estimation of the diffusion time (See *<u>micapipe-supplementary</u>* for details). Group-level gradients were constructed from the average of subject-level cortical matrices. For MPC, FC, and GD, matrices were thresholded row-wise to retain the top 20% edges (see ‘*<u>Building gradients</u>*’ in the documentation, **Figure S3** for an example at the MICs dataset). SC matrices were log-transformed to reduce connectivity strength variance, but not thresholded. Moreover, left and right hemispheres were analyzed separately for SC, given limitations of diffusion tractography in mapping inter-hemispheric fibers. Hemispheres were also analyzed separately for GD gradients, as the surface-based measure of geodesic distance used here is computed on distinct hemisphere surface spheres.. The For each subject, we aligned the first gradient using Procrustes rotations to the group-level gradient for each modality, and computed correlations as a measure of inter-subject consistency.

##### Graph features

Graph measurements were computed using the igraph R package (<u>igraph.org/r</u>). We focused on three widely used graph-theoretical parameters, node strength, characteristic path length, and clustering coefficient (Rubinov and Sporns, 2010). We computed the clustering-coefficient as a measure of segregation, which provides information about the level of local connections in a network.

The characteristic path length quantified network integration with short path lengths indicating globally efficient networks. Dijkstra’s algorithm was used to calculate the inverse distance matrix (Dijkstra, 1959) and infinite path lengths were replaced with the maximum finite length (Van den Heuvel et al., 2008). Finally, we calculated strength to characterize the relevance of the individual nodes. FC strength was calculated only with positive values. Using the same thresholding as for the diffusion map embedding, GD, MPC and FC matrices were thresholded to retain the top 20% of the edges, and SC was analyzed using the un-thresholded, weighted networks.

#### Inter-datasets similarity

To assess stability across datasets, we computed Spearman’s correlation coefficients between the group-level measures of each pair of datasets for each MRI modality (**Figure 4**).

### 4.6 Version control and containers

*micapipe* is executable via a Docker container, and we provide information on how to convert it to a singularity image either via directly pulling from *<u>dockerhub</u>* or converting a local image (Kurtzer et al., 2017). Each new version of *micapipe* is uploaded and tagged, and changes are documented. The current release version is v.0.1.2. Our goal is to maintain continuous integration. Additionally, our pipeline has adopted the standards of BIDS-Apps (Gorgolewski et al., 2017) and of the center for reproducible neuroimaging computation (Robert et al., 2016).

## 5. Acknowledgements

RRC received support from the Fonds de la Recherche du Québec – Santé (FRQ-S). JR received support from the Canadian Open Neuroscience Platform (CONP) and Canadian Institute of Health Research (CIHR). SL acknowledges funding from FRQ-S, CIHR, and the Richard and Ann Sievers Neuroscience Award. RVdW received support from the Savoy Foundation and the Richard and Ann Sievers Neuroscience Award. OB received support from the Healthy Brains for Healthy Lives (HBHL) program and the Transforming Autism Care Consortium (TACC). BP was supported by the National Research Foundation of Korea (NRF-2021R1F1A1052303), Institute for Information and Communications Technology Planning and Evaluation (IITP) funded by the Korea Government (MSIT) (2020-0-01389, Artificial Intelligence Convergence Research Center, Inha University; 2021-0-02068, Artificial Intelligence Innovation Hub), and Institute for Basic Science (IBS-R015-D1). LC acknowledges support from CONACYT (181508, 1782) and from UNAM-DGAPA (IB201712, IG200117, IN204720). BCB acknowledges support from CIHR (FDN-154298, PJT-174995), SickKids Foundation (NI17-039), Natural Sciences and Engineering Research Council (NSERC; Discovery-1304413), Azrieli Center for Autism Research of the Montreal Neurological Institute (ACAR), BrainCanada, FRQ-S, the Helmholtz International BigBrain Analytics and Learning Laboratory (Hiball), and the Canada Research Chairs Program (CRC). P.H. was supported in parts by funding from the Canada First Research Excellence Fund, awarded to McGill University for the Healthy Brains for Healthy Lives initiative, the National Institutes of Health (NIH) NIH-NIBIB P41 EB019936 (ReproNim), the National Institute Of Mental Health of the NIH under Award Number R01MH096906, a research scholar award from Brain Canada, in partnership with Health Canada, for the CONP initiative, as well as an Excellence Scholarship from Unifying Neuroscience and Artificial Intelligence - Québec.

## 7. Supplementary figures and tables

**Figure S1.**
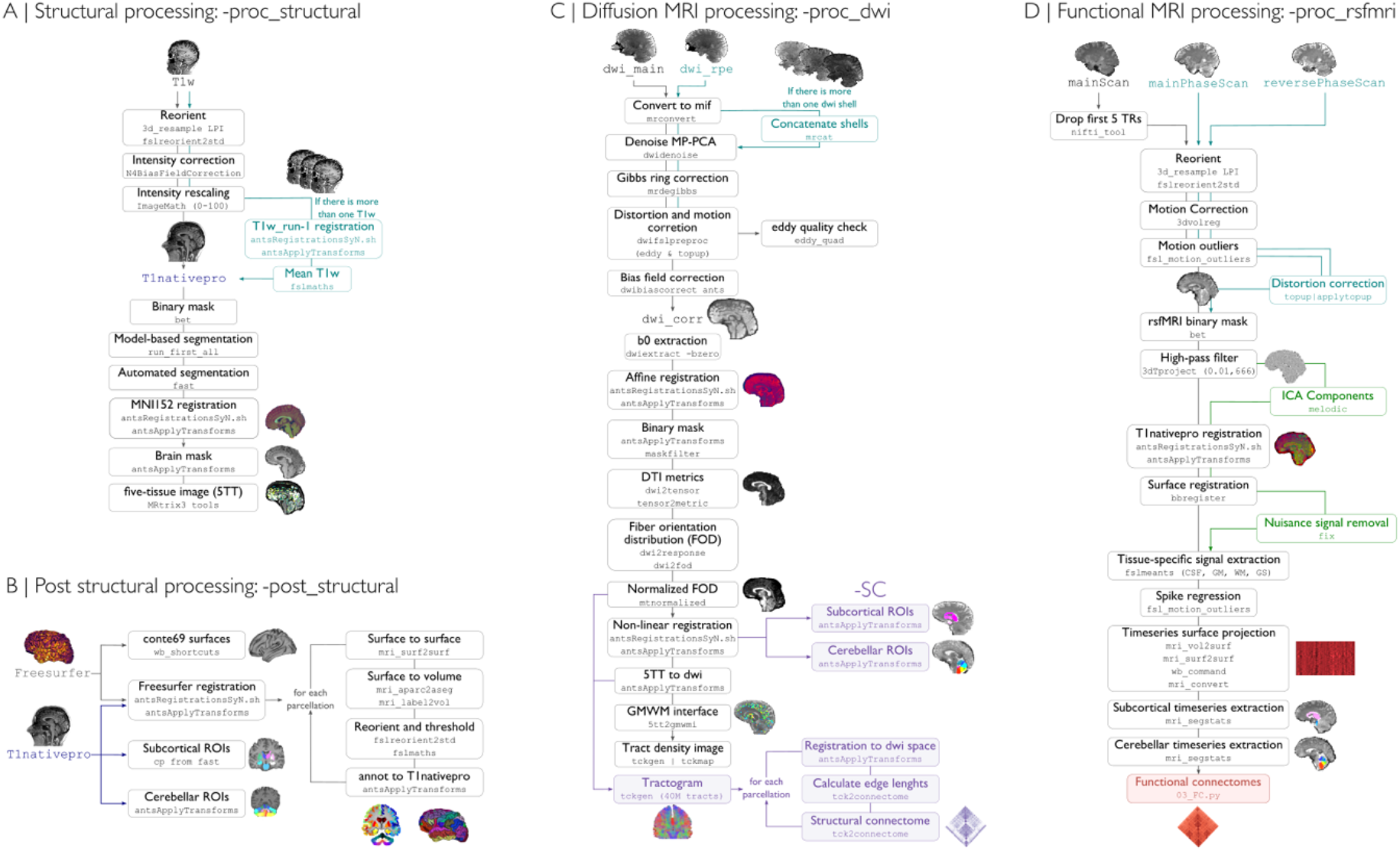
Core processing modules. **A)** The <u>-proc_structural</u> module inputs a T1-weighted image, and generates a native space processed image (nativepro). This image will be used for registration purposes in all following steps. **B)** After FreeSurfer is run and T1nativepro is generated, the <u>-post_structural</u> module registers native FreeSurfer surfaces to a standard template, in addition to mapping all cortical parcellation schemes to the subject’s native surface space and nativepro space. **C)** The <u>-proc_dwi</u> processes the native DWI data, and estimates a structural connectome (via <u>-SC</u>). We apply iFOD2 for this purpose, a probabilistic tractography algorithm. **D)** The <u>-proc_rsfmri</u> module performs all pre-processing of the rs-fMRI scans, in preparation for the construction of functional connectomes.

**Figure S2.**
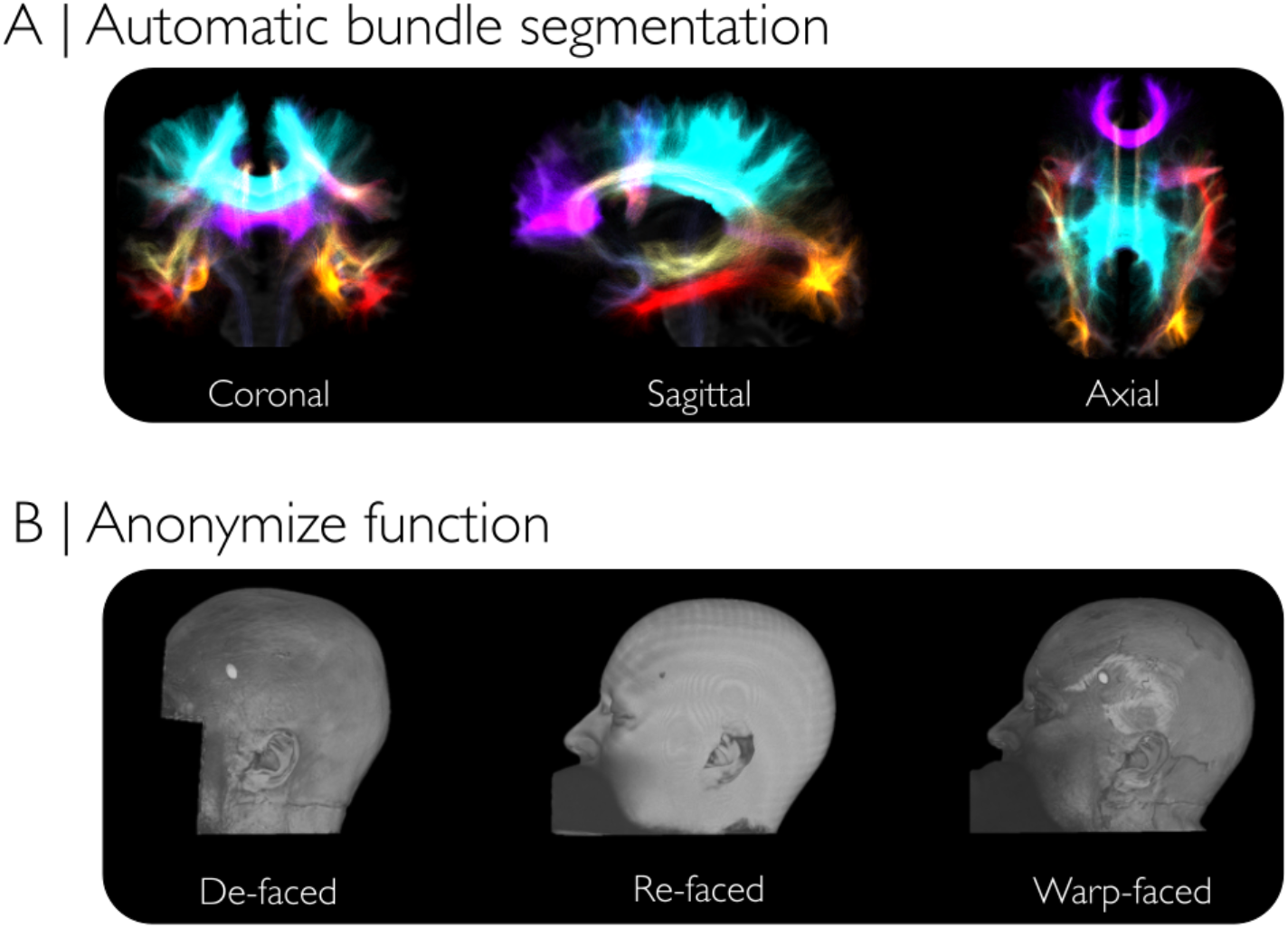
Additional features. **A.** Automatic bundle segmentation generating 35 virtually dissected white matter bundles. Segmented tracts obtained from whole-brain tractography generated in the -SC module. **B.** Anonymizing function for the anatomical images from the BIDS directory. Three different methods are available for defacing/refacing.

**Figure S3.**
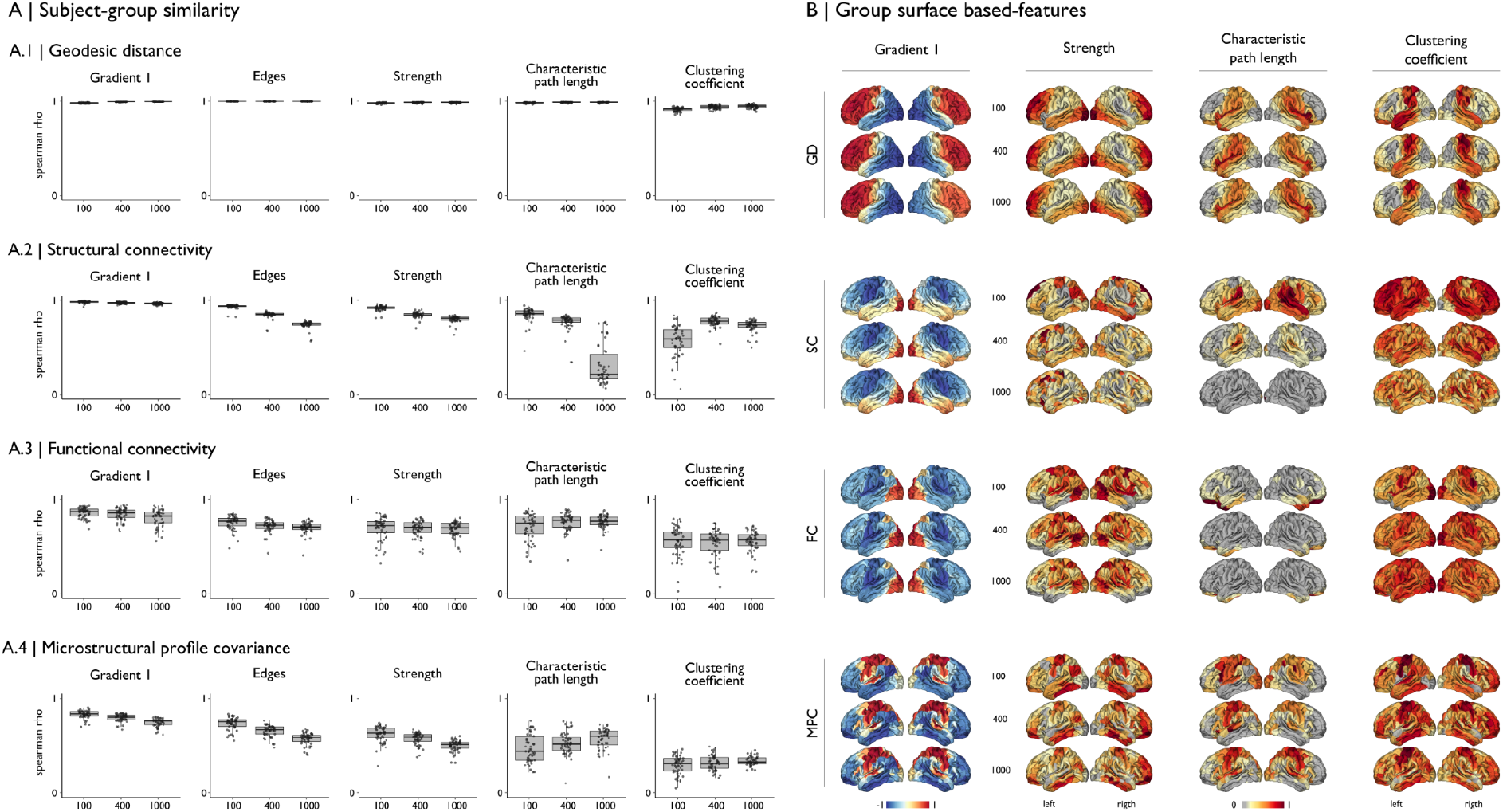
Within-dataset consistency. A) Subject-group consistency for each modality and measurement in the MICs dataset. Each point in the boxplots corresponds to a Spearman’s rho between each subject and the group average measurement, derived from the Schäfer-100,−400 and 1000 parcellations. B) Group average feature maps from the MICs dataset.

**Table S1.** Evaluation datasets. For further details about the code used to run these datasets see *<u>micapipe.readthedocs/databases</u>*. The T1w/T2w or Flair, the weighted image was calculated based on these structural acquisitions. FOV: field of view, TE: repetition time, TE: echo time, T1w: T1-weighted.

| Database | Field | Subjects /sessions | T1w | DWI main | DWI rpe | rs-fMRI | rs-fMRI rpe | Quantitative Image |
| --- | --- | --- | --- | --- | --- | --- | --- | --- |
| *CamCan* | 3T Siemens TIM Trio | 130/1 | 1mm<sup>3</sup> isotropic, FOV=192x256x256 | 2mm<sup>3</sup> isotropic, FOV=96x69x66, 30-b1000, and 30-b2000, 3-b0 | No | 3x3x4.44mm<sup>3</sup>, FOV=64x64x32x246 TE=0.03, TR=1.97 | No | MTR: 1.6mm<sup>3</sup> isotropic, FOV=104x128x128 |
| *SUDMEX* | 3T Philips Ingenia | 145/1 | 1mm<sup>3</sup> isotropic, FOV=180x240x240 | 2mm<sup>3</sup> isotropic, FOV=112x112x50, 32-b1000, and 96-b3000, 8-b0 | Yes | 3mm<sup>3</sup> isotropic, FOV=80x80x36x300 TE=0.03, TR=2 | Yes |  |
| *EpiC* | 3T Philips Achieva TX | 78/1 | 1mm<sup>3</sup> isotropic, FOV=176x256x265 | 2mm<sup>3</sup> isotropic, FOV=128x128x50, 60-b2000 and 1-b0 | Yes | 1.8x1.8x3mm<sup>3</sup>, FOV=128x128x33x200 TE=0.03, TR=2 | No | T1w/Flair |
| *EpiC longitudinal* | 3T Philips Achieva TX | 33/1 | 1mm<sup>3</sup> isotropic, FOV=176x256x265 | 2mm<sup>3</sup> isotropic, FOV=128x128x66, 60-b1000, 60-b2000 and 2-b0 | Yes | 1.8x1.8x3mm<sup>3</sup>, FOV=128x128x33x200 TE=0.03, TR=2 |  | T1w/Flair |
| *MSC* | 3T Siemens TRIO | 9/12 | 0.8mm<sup>3</sup> isotropic, FOV=224x256x256 | No | No | 4mm<sup>3</sup> isotropic, FOV=64x64x36x818 TE=0.027, TR=2.2 | No | T1w/T2w |
| *MICs* | 3T Siemens Magnetom prisma fit | 50/1 | 0.8mm<sup>3</sup> isotropic, FOV=224x320x320 | 1.6mm<sup>3</sup> isotropic, FOV=140x140x93, 10-b300, 40-b1000, 90-b200 3-b0 | Yes | 3mm<sup>3</sup> isotropic, FOV=80x80x48x700 TE=0.03, TR=0.6 | Yes | qT1w: 0.8mm<sup>3</sup> isotropic, FOV=240x320x320, TE=0.0029, TR=5 |
| *Audiopath* | 7T Siemens Magnetom | 10/1 | 0.6mm<sup>3</sup> isotropic, FOV=256x384x384 | 1mm<sup>3</sup> isotropic, FOV=200x200x132, 66-b1000, 66-b2000, 66-b3000, 33-b0 | All | data not processed (cropped) | Yes | T1w/T2w?? |

**Table S2.** Processing times (shown as the mean ± SD across subjects in minutes). ***pp***: preprocessed, ***np***: not processed, No data available. The Morphology module took about 1 minute to process in all the databases. Multi-threaded processing is available only for ANTs and workbench functions.

| Module/Dataset | MICs | EpiC | EpiC<br>longitudinal | Cam-Can | MSC | SUDMEX | Audiopath |
| --- | --- | --- | --- | --- | --- | --- | --- |
| proc_structural | 88±17 | 76±9 | 73±10 | 70±6 | 76±2 | 66±4 | 92±3 |
| post_structural | 125±14 | 44±10 | 46±14 | 51±7 | 135±20 | 44±4 | 227±22 |
| proc_freesurfer | 961±205 | *pp* | *pp* | *pp* | 815±88 | *pp* | 1154±104 |
| GD | 159±21 | 119±40 | 96±12 | 105±12 | 162±15 | 100±8 | 242±36 |
| proc_dwi | 246±37 | 63±15 | 114±15 | 41±5 | -- | 99±12 | 1104±274 |
| SC | 906±427 | 1364±498 | 1471±538 | 1086±387 | -- | 1652±776 | 910±399 |
| proc_rsfmri | 101±8 | 22±3 | 22±2 | 15±4 | 79±54 | 28±7 | np |
| MPC | 7±1 | 5±1 | 5±1 | 6±1 | 8±1 | -- | 9±2 |
| Mean total time<br>by subject<br>(minutes) | 2593±730 | 1693±576 | 1827±592 | 1374±422 | 1275±180 | 1989±811 | 3738±840 |

**Table S3.** Mean total size of the main directories by subject and database. The size of the directories is shown as mean and SD in Megabytes. The databases with modules that were not processed are shown as 0.

| Dataset | MICs | EpiC | EpiC longitudinal | Cam-Can | MSC | SUDMEX | Audiopath |
| --- | --- | --- | --- | --- | --- | --- | --- |
| anat | 401 ± 16 | 779 ± 424 | 326 ± 72 | 293 ± 39 | 89 ± 181 | 219 ± 13 | 591 ± 68 |
| dwi | 3447 ± 10 | 2731 ± 856 | 1660 ± 188 | 726 ± 105 | 0 | 1126 ± 374 | 18480 ± 1754 |
| func | 8647 ± 498 | 2221 ± 633 | 2034 ± 945 | 1949 ± 297 | 4867 ± 3816 | 3370 ± 854 | 0 |
| xfm | 1474 ± 69 | 1123 ± 52 | 1114 ± 135 | 1014 ± 30 | 273 ± 485 | 901 ± 25 | 3840 ± 374 |
| QC | 103 ± 6 | 37 ± 20 | 57 ± 37 | 46 ± 13 | 0 | 43 ± 16 | 106 ± 41 |
| Total | 14072 ± 599 | 6891 ± 1985 | 5191 ± 1377 | 4028 ± 484 | 5229 ± 4482 | 5659 ± 1282 | 23017 ± 2237 |

**Table S4.** rs-fMRI processing. Parameters used to process the rs-fMRI data. For details see <u>micapipe.readthedocs/databases</u>.

| Database | Topup | Melodic | FIX | CSF/WM<br>regression | Non-linear<br>registration | GS<br>regression |
| --- | --- | --- | --- | --- | --- | --- |
| *CamCan* | *No* | *No* | *No* | **Yes** | **Yes** | *No* |
| *SUDMEX* | **Yes** | *No* | *No* | **Yes** | **Yes** | *No* |
| *EpiC* | *No* | *No* | *No* | **Yes** | **Yes** | *No* |
| *EpiC*<br>*longitudinal* | *No* | *No* | *No* | **Yes** | **Yes** | *No* |
| *MSC* | *No* | *No* | *No* | **Yes** | **Yes** | *No* |
| *MICs* | **Yes** | **Yes** | **Yes** | *No* | **Yes** | *No* |

